# Weakly supervised contrastive learning infers molecular subtypes and recurrence of breast cancer from unannotated pathology images

**DOI:** 10.1101/2023.04.13.536813

**Authors:** Hui Liu, Yang Zhang, Aichun Zhu, Zhiqiang Sun, Judong Luo

**Affiliations:** College of Computer and Information Engineering, Nanjing Tech University, Nanjing, 211816, Jiangsu, China.; Department of Radiotherapy, The Affiliated Changzhou Second People’s Hospital of Nanjing Medical University, City, 213110, Jiangsu, China.

**Keywords:** Contrastive learning, pathology images, weakly supervised learning, molecular subtyping, cancer recurrence, spatial transcriptome

## Abstract

The deep learning-powered computational pathology has led to significant improvements in the speed and precise of tumor diagnosis,, while also exhibiting substantial potential to infer genetic mutations and gene expression levels. However,current studies remain limited in predicting molecular subtypes and recurrence risk in breast cancer. In this paper, we proposed a weakly supervised contrastive learning framework to address this challenge. Our framework first performed contrastive learning pretraining on large-scale unlabeled patches tiled from whole slide images (WSIs) to extract patch-level features. The gated attention mechanism was leveraged to aggregate patch-level features to produce slide feature that was then applied to various downstream tasks. To confirm the effectiveness of the proposed method, we have conducted extensive experiments on four independent cohorts of breast cancer. For gene expression prediction task, rather than one model per gene, we adopted multitask learning to infer the expression levels of 21 recurrence-related genes, and achieved remarkable performance and generalizability that were validated on an external cohort. Particularly, the predictive power to infer molecular subtypes and recurrence events was strongly validated by cross-cohort experiments. In addition, the learned patch-level attention scores enabled us to generate heatmaps that were highly consistent with pathologist annotations and spatial transcriptomic data. These findings demonstrated that our model effectively established the high-order genotype-phenotype associations, thereby enhances the potential of digital pathology in clinical applications.

## 1 Introduction

Breast cancer is one of the most prevalent and lethal malignant neoplasms among women. According to a recent report, there were 2.6 million cases of breast cancer in 2020 [1]. Breast cancer often exhibits considerable tumor heterogeneity, with marked variability in molecular subtypes, treatment outcomes, and prognosis among individual patients. At present, genetic testing is the standard methodology for molecular subtyping and pre-treatment efficacy assessment. However, the high cost and long turnaround time of genetic testing impede its widespread application. Consequently, there is an pressing demand in clinical practice for the development of a new, cost-effective and scalable alternative approach.

With the advent of whole-slide imaging technology and the growing emphasis on precision medicine, AI-powered digital pathology has facilitated the extraction of histological features beyond human visual perception ability. Digital pathology has emerged as an efficacious approach to inferring genetic alteration and molecular signatures. For example, Noorbakhsh et al. developed a convolutional neural network (CNN) model that effectively differentiated between tumor/normal tissue and cancer subtypes among 19 types of cancer[2]. Wang et al. proposed a deep learning model to study the tumor microenvironment using pathological images and uncovered that cellular spatial organization is predictive of gene expression profiles and patient survival[3]. Wang et al. developed a deep learning-based histological grading model (DeepGrade) based on whole-slide images (WSI) to improve risk stratification for NHG 2 breast cancer patients and validated the effectiveness of their model [4]. Kather et al. employed a deep residual network to directly predict microsatellite instability from pathological images[5]. Jain et al. utilized deep learning to predict tumor mutation burden from pathological images [6]. Saltz et al. leveraged deep learning to reveal the tumor immune microenvironment and lymphocyte infiltration[7].

The aforementioned studies illustrate the remarkable potential of deep learning in the realm of digital pathology image analysis. However, this approach is also confronted with the challenge of a paucity of pixel-level manual annotation, as it is both costly and time-consuming, due to experienced pathologists are exceedingly scarce and busy. To address this challenge, some researchers have turned to semi-supervised and weakly supervised learning techniques. For example, Mahmood et al. developed the CLAM model, which utilizes weakly supervised learning and attention mechanisms to classify subtypes of renal cell carcinoma and non-small cell lung cancer using pathological images[8]. Zhang et al. devised a dual-layer feature distillation multi-instance learning model that demonstrated impressive performance on the CAMELYON-16 and TCGA lung cancer datasets[9]. Wu et al. harnessed the power of consistency regularization and entropy minimization to develop an efficacious semi-supervised kernel image segmentation algorithm[10].

In recent years, the rapid advancement of multi-omics technology has provided novel insights into the tumor molecular subtypes, biomarkers of prognosis and drug response. For instance, gene expression profiling has facilitated the establishment of molecular subtypes of breast cancer[11]. A 12-gene expression signature has been developed as an independent indicator of recurrence for stage II colon cancer patients [12]. Furthermore, gene signatures have been employed to establish recurrence risk models for early-stage breast cancer patients [13][14][15]. However, due to its high cost, molecular diagnosis based on gene sequencing cannot be applied widespread in clinical practice. Consequently, researchers have endeavored to infer gene expression profiles based on high-resolution pathological images. For example, Kather et al. demonstrated that pan-cancer genetic alterations could be inferred from pathological images [16]. Schmauch et al. proposed the HE2RNA model to predict RNA-seq expression from pathological images and identify tumors with microsatellite instability [17]. Huang et al. developed the HistCode model to predict differential expression of tumor driver genes [18].

This paper proposed a deep learning framework based on weakly supervised contrastive learning, with the aim of establishing the links between molecular characteristics and histopathological morphology. We first splited WSIs to millions of small patches, and then performed contrastive learning pretraining on the large-scale unlabeled patches to extract patch-level embeddings. For weakly supervised learning without annotation, the patch-level embeddings were aggregated using a gated attention mechanism to obtain slide-level feature, which is subsequently used in downstream whole-slide-level tasks. To validate the effectiveness of our method, we leveraged the computational pathological feature to infer gene expressions, molecular subtypes, treatment responses and clinical outcome. The extensive experiments demonstrated that our method can accurately identify tumor regions using only weakly supervised labels and exhibits exceptional performance in predicting breast cancer-related gene expression levels, as well as multiple clinically relevant tasks. Furthermore, spatial heatmaps generated based on the patch-level attention scores learned in tumor diagnosis and gene expression inference tasks exhibit high consistency with pathologist annotations and spatial transcriptomic data. This indicates that the deep learning-based model effectively establishes the genotype-phenotype links, greatly enhancing the potential of digital pathology in clinical applications.

## 2 Results

### 2.1 Weakly supervised contrastive learning framework

Our study proposed a computational pathology framework that integrated selfsupervised contrastive learning and attention-based multiple-instance aggregation to enable multi-task weakly supervised tasks based on the whole-slide images (WSIs). The framework comprised three stages (Figure 1). In the first stage, super high-resolution whole slide images were split into small 256×256px patches. The patches without enough detectable foreground tissue by redefined area threshold were discarded. The second stage run contrastive learning-based pretraining on the large-scale unannotated patches to obtain informative patch-level features. In the three stage, the gated attention mechanism was employed to aggregate patch-level features into slide-level representation. For each downstream tasks, the parameters of the image encoder were frozen to learn the attention score of each patch based on weakly supervisory signals.

**Fig. 1:**
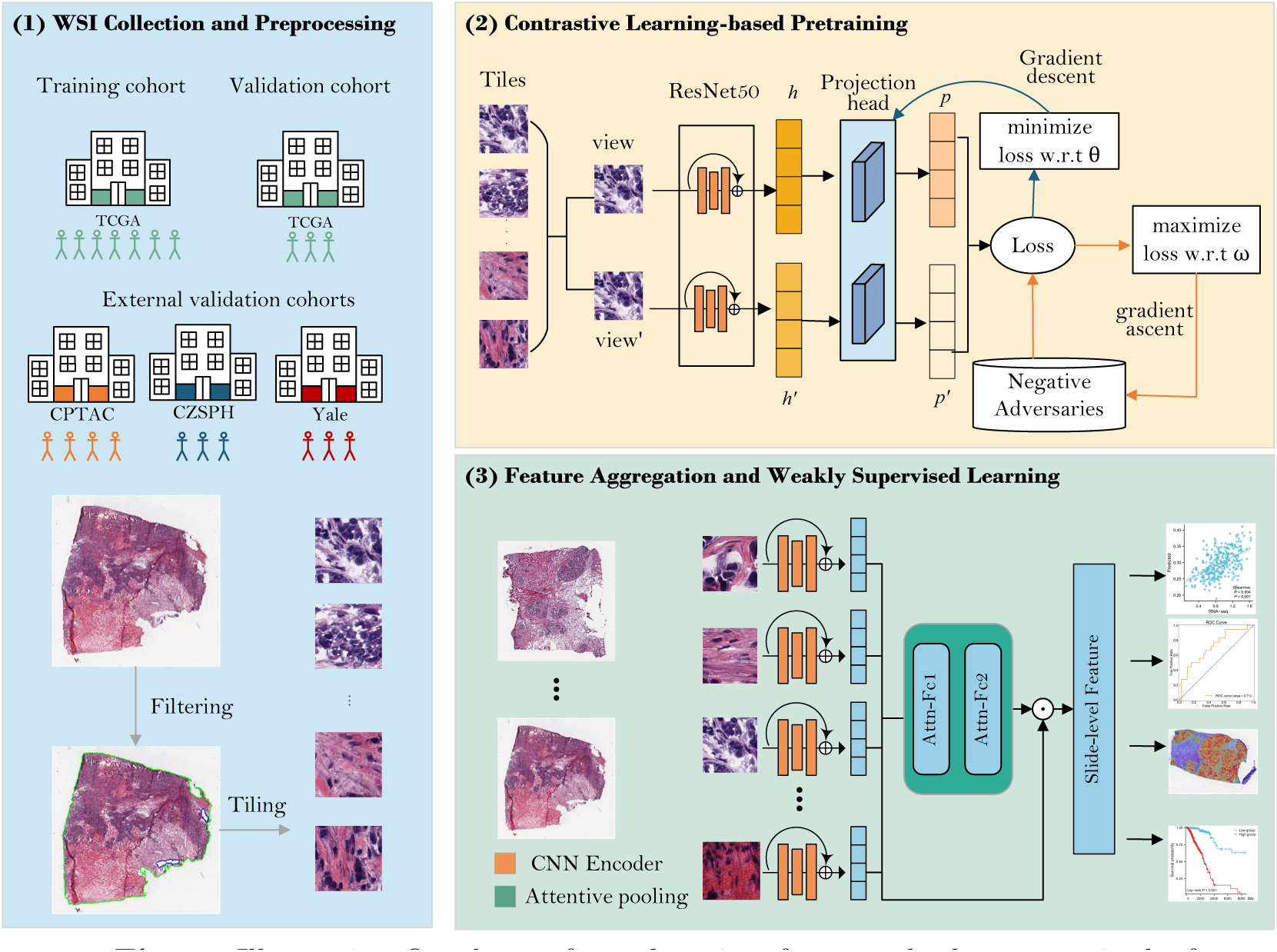
Illustrative flowchart of our learning framework that comprised of three steps: (1) H&E slide images of four breast cancer cohorts were collected from different resources. After filtration and segmentation, each WSI was tiled into numbers of patches. (2) The Adversarial Contrastive learning (AdCo) was leveraged to learn latent representations on large-scale unlabeled patches. AdCo algorithm uses the contrastive learning and adversarial training strategy to learn low-dimensional but informative embedding for an input image. (3) During training and inference, an attention network is used to aggregate the patch-level features into sliding-level representations, which were applied to multiple clinically relevant tasks. The attention scores can be utilized to build heatmaps for identifying ROIs and interpret the important morphology for different downstream tasks.

Unlike previous studies, we utilized self-supervised contrastive learning pretraining to extract features from millions of patches. Contrastive learning seeks to minimize the distance between original and augmented images while simultaneously maximizing their distance from negative samplses within a minibatch. This approach encourages the encoder to capture the essential feature of pathology images, resulting in more expressive embedding representations. Our previous work [18] and experiments conducted in this study (see Subsection 2.8) strongly supported that pretraining based on contrastive learning significantly enhances the predictive ability in downstream tasks.

Notably, our approach did not require pixel-level or region-of-interest (ROI) annotations. In fact, our model considered all patches in the tissue regions of a WSI, and learned an attention score to each patch that conveyed its contribution or importance to the collective slide-level representation for a specific downstream task. This interpretation of the attention score was reflected in the slide-level aggregation rule of attention-based pooling, which computed the slide-level representation as the average of all patches in the slide weighted by their respective attention score. The patch-level attention score enabled spatial localization of tumor subregions and gene expressions by spatial deconvolution of patches to original slide (see Subsection 2.9).

### 2.2 Accurate tumor diagnosis and tumor region localization using weakly supervisory signals

We first evaluated the proposed model on the TCGA-BRCA cohort by conducting a tumor diagnosis task. This classification task utilized only samples from fresh frozen tissue sections, comprising a total of 1,807 pathological slides, including 1,566 tumor tissues and 241 normal tissues. For objective performance assessment of our model, we randomly allocated 30% WSIs as an independent test set, and the remainder as training set, with the hyperparameters fine-tuned by 3-fold cross-validation. Given that we only had slide-level labels, we employed weakly supervised learning for tumor diagnosis and attempted to localize tumor regions. Specifically, each WSI was treated as a set composed of patches. The patch features obtained in the pretraining stage were aggregated into slide features through attentive pooling and subsequently used to predict whether or not the slide contained tumor tissue. On the TCGA-BRCA cohort, our method achieved an accuracy of 0.99 and an ROC-AUC value of 0.99. Figure 2(a-b) displayed the ROC curve and confusion matrix on the test set. This result strongly supported that our model can accurately learn to distinguish whether a WSI contains tumor tissue using only weak supervisory signals. By performing UMAP dimensionality reduction on the latent representation of the slides, we discovered that tumor and non-tumor slides are significantly separated in the embedding space.

**Fig. 2:**
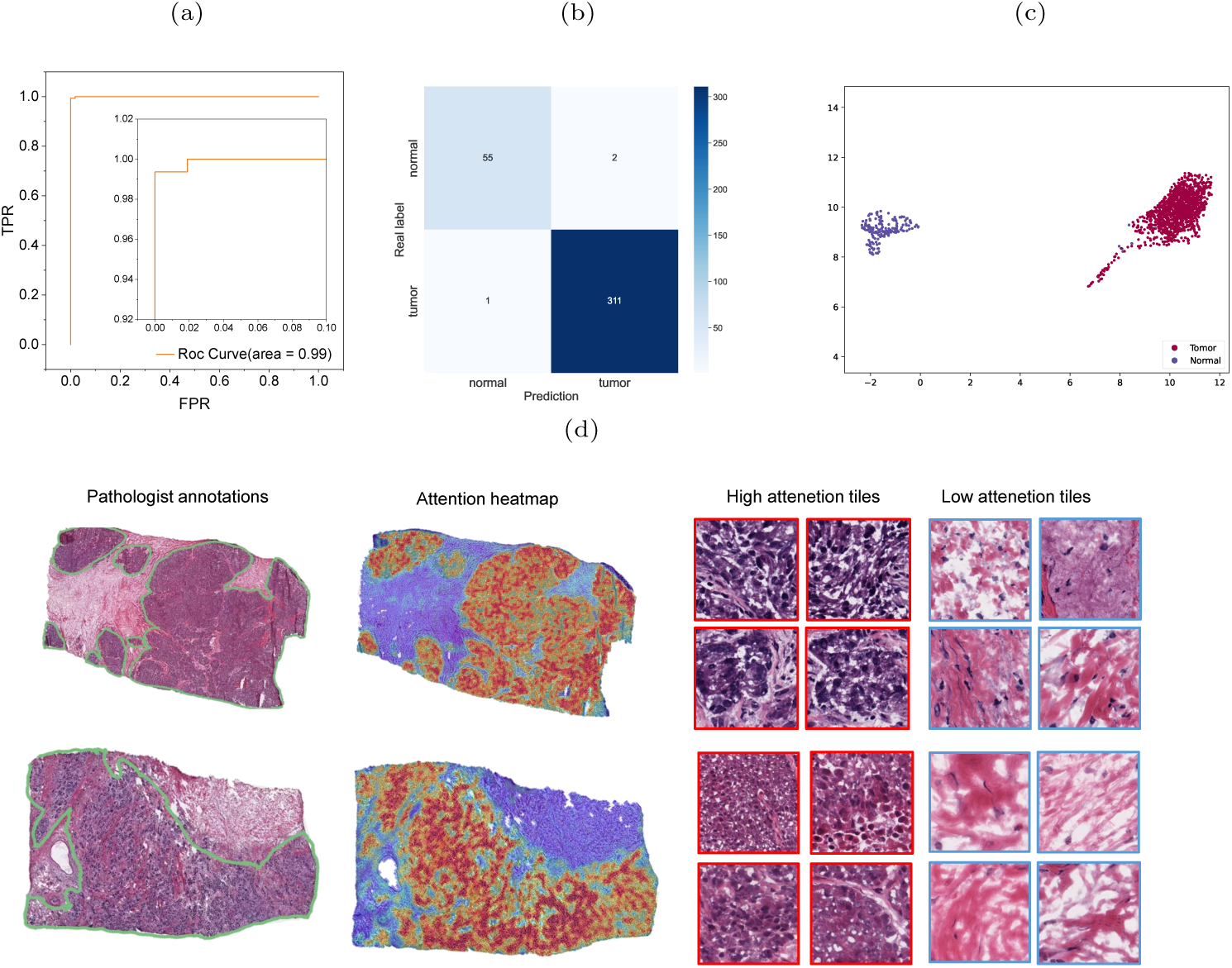
Performance evaluation for tumor diagnosis task based on computational pathological features. (a-b) ROC curve and confusion matrix achieved by our method on the leave-out test set of TCGA-BRCA cohort. (c) UMAP visualization of the learned slide-level representations showed that tumor and normal slides separated remarkably. (d) Spatial localization of the tumor regions uncovered by the learned patch-level attention scores. Left column showed the tumor and necrosis regions annotated by an experienced pathologist. The middle column displayed the heatmaps generated by spatially deconvoluting each patch to original slide and colored each patch using its learned attention score. The right column showed some representative patches assigned high and low attention scores.

Since the slide features for tumor diagnosis are obtained by aggregating the features of patches through an attention mechanism, patches with high attention scores imply tumor regions. To assess whether our model can effectively locate tumor regions, we normalized the patch attention scores learned in the tumor diagnosis task and spatially deconvoluted each patch to generate heatmaps. Concurrently, we invited a pathologist with 10 years of experience to manually annotate the tumor regions. As depicted in Figure 2(d), the high-attention regions exhibited high consistency with the tumor areas annotated by the pathologist. Several selected patches corresponding to high and low attention patches displayed evident differences in tissue morphology.

### 2.3 Multitask learning predicted the gene expression profiles relevant to breast cancer recurrence

To confirm that our model learned real and informative features from pathological images, we constructed an MLP model to predict the expression levels of 21 genes that have been confirmed to be closely related to breast cancer recurrence in previous studies [13]. Unlike existing studies that built one model per gene [19], we adopted multi-task learning to quantitatively predict the expression levels of 21 genes simultaneously. For this purpose, we downloaded RNA-seq data (FPKM-UQ) with matched WSI from TCGA-BRCA cohort, and obtained a total of 1,572 paired samples. We randomly selected 20% used as an independent test set and the remaining as training set, with hyperparameters tuned by 4-fold cross-validation. For this regression task, we used the correlation coefficient as the performance evaluation metric. Our method achieved an average correlation coefficient of 0.36, and the histogram of predicted correlation coefficients for 21 genes was shown in Figure 3(a). Further, we examined the top 10 well-predicted genes and found that most of them were related to cell proliferation. For example, MKI67 (r=0.504) is a wellknown cell proliferation marker expressed in both tumor and non-tumor cells. Its overexpression has been found to be associated with tumor growth, histological staging, and tumor recurrence[20–22]. ESR1 (r=0.555) mutation is an important cause of endocrine resistance and acts as a clinical biomarker for metastatic hormone receptor-positive breast cancer[23]. AURKA (r=0.58) belongs to the serine/threonine kinase family and its activation is essential for regulating cell division through mitosis and has been shown to be associated with prognosis in ER-positive, EBBR2-negative and TN breast cancer subtypes [24]. ERRB2 (r=0.252) overexpression/amplification is associated with malignant tumors and poor prognosis in breast cancer and is an important therapeutic target for breast cancer[25].

**Fig. 3:**
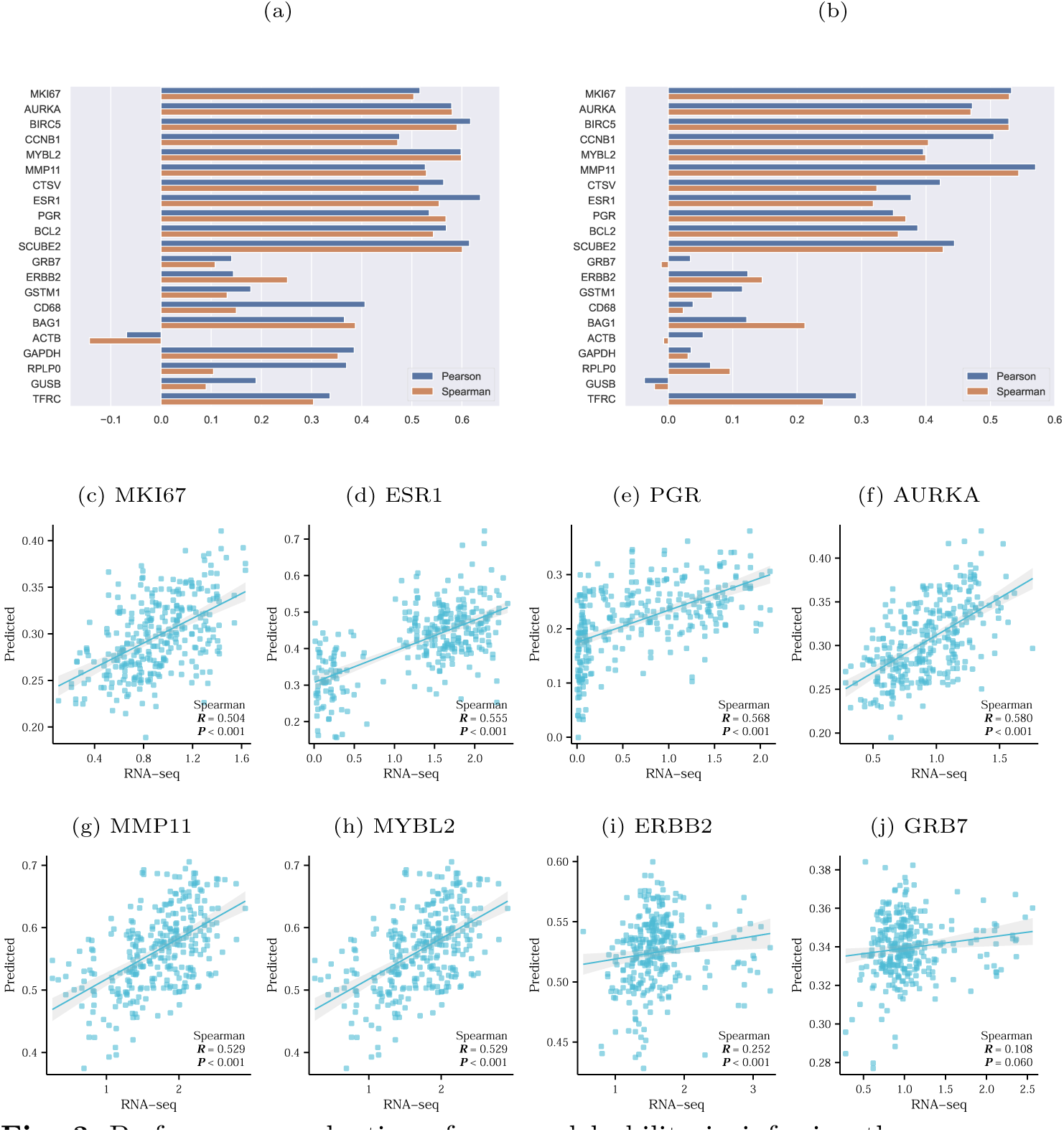
Performance evaluation of our model ability in inferring the expression levels of 21 recurrence-related gene using the computational pathological features. (a) Histogram of the correlation coefficients achieved on the leaveout test set of TCGA-BRCA cohort. (b) Histogram of correlation coefficients achieved by our model trained on TCGA-BRCA cohort and evaluated on the external CPTAC-BRCA cohort. (c-j) Scatter plots of the predicted and actual expression levels of eight genes closely associated with breast cancer. For each gene, the Spearman coefficient and *p*-value were also shown.

### 2.4 External cohort validated the generalizability of gene expression level prediction

Unlike studies that built one prediction model per gene, our multi-task learning model demonstrated superior robustness. To further evaluate the generalizability of our model in predicting gene expression levels, we employed an independent external cohort from the CPTAC-BRCA project, consisting of 97 samples with matched pathological images and RNA-seq data. Notably, instead of training and testing on the external dataset, we completed model training on the TCGA-BRCA cohort and subsequently tested it on the CPTAC cohort to confirm that our model captured authentic features from a large number of patches. Our model achieved an average correlation coefficient of 0.26, with the highest correlation coefficient for the MMP11 gene reaching 0.54 (p*<*0.01), as illustrated in Figure 3(b). These results indicated that deep learning-based gene expression prediction from pathological images can be generalized to external populations. The experimental results demonstrated that our method has made a significant progress towards practical clinical application.

### 2.5 Cross-cohort validation demonstrated generalizability in predicting breast cancer recurrence

We further applied computational pathological features to predict the five-year recurrence of breast cancer. We evaluated the performance of our model on two datasets: the TCGA-BRCA cohort (n=272) and the CZSPH cohort (n=91). A case was labeled as 1 if it had recurrence, metastasis, or a new tumor event within five years, and 0 otherwise. We applied three-fold cross-validation and AUC as the evaluation metrics. Figure 4(a) showed that our model achieved an AUC of 0.71 on the independent test set randomly sampled from the TCGA-BRCA cohort. To further test the generalizability of our model, we predicted patient recurrence events in the CZSPH cohort using the model trained on the TCGA-BRCA cohort. We considered this cross-cohort prediction experiment as a strong validation of our model robustness. Figure 4(b) illustrated that the model attained an AUC of 0.69. These results indicated that our model has high effectiveness and generalizability in predicting breast cancer recurrence.

**Fig. 4:**
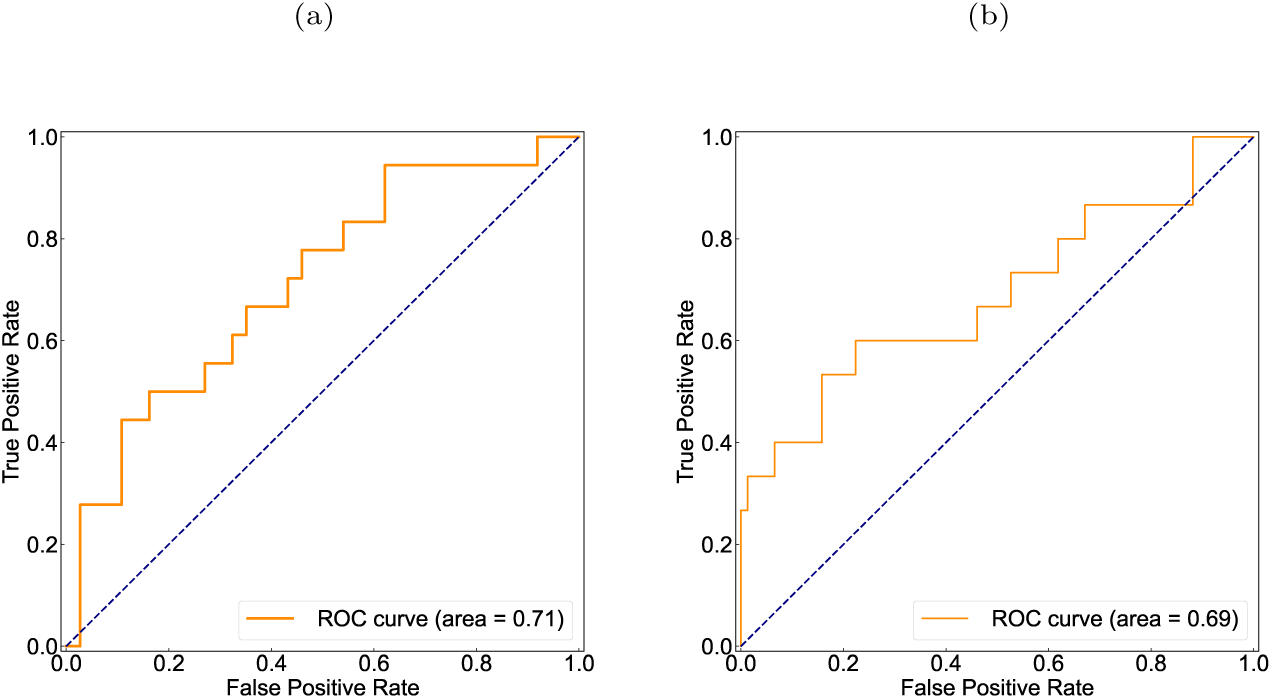
Performance evaluation in prediction of five-year recurrence events in breast cancer based on the extracted pathological features by our model. (a) ROC curve and AUC value achieved by our model on the leave-out test set of TCGA-BRCA cohort. (b) ROC curve and AUC value achieved by our model trained on TCGA-BRCA cohort and evaluated on the independent external CZSPH cohort.

### 2.6 Accurate prediction of breast cancer molecular subtypes

PAM50 is a 50-gene signature that classifies breast cancer into five molecular intrinsic subtypes: Luminal A, Luminal B, HER2-enriched, Basal-like, and Normal-like [11]. Each subtype has distinct molecular characteristics, prognosis, clinical behavior, and treatment response [26, 27]. For instance, luminal A and B subtypes are associated with relatively favorable outcomes and are mainly treated with surgery, chemotherapy, and endocrine therapy. Basal-like subtype is characterized by poor survival and lack of effective treatment options. Her2-enriched subtype requires targeted drugs or radiotherapy before surgical resection. In this study, we developed a model to predict PAM50 sub-types using pathological image features. Figure 5(a-b) showed that our model achieved high performance in predicting all four subtypes, with an average ROC-AUC value of 0.88 and PR-AUC value of 0.73. Notably, the AUC for the basal-like subtype reached 0.96. Our model can provide clinically actionable results by accurately predicting PAM50 subtypes using routine pathological images, especially for the basal-like subtype, which is otherwise costly and inaccessible in clinical practice.

**Fig. 5:**
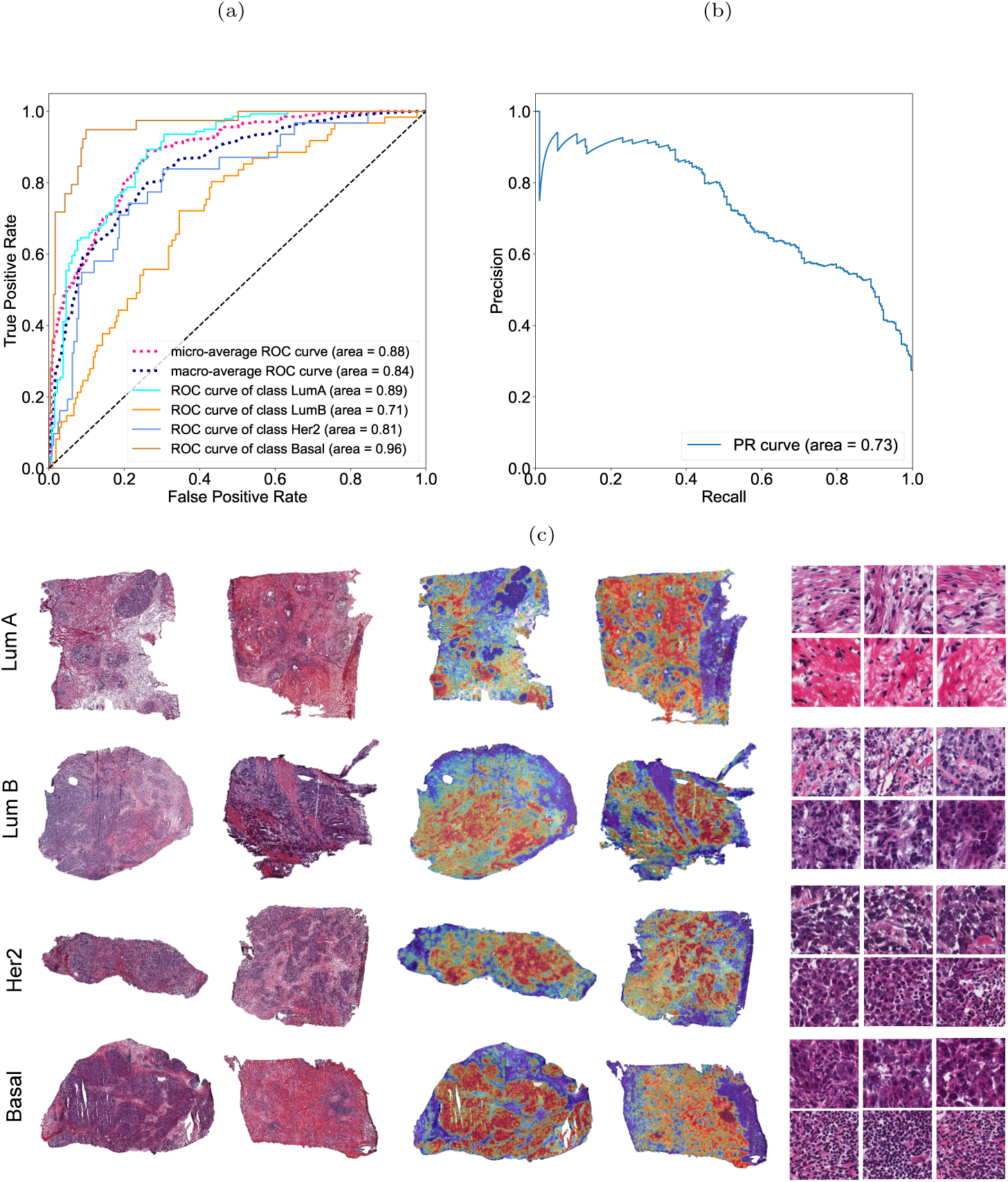
Evaluation of our model performance in predicting molecular subtypes of breast cancer, and visualization of attention heatmaps corresponding to each molecular subtype. (a) ROC curves and AUC values achieved by our model on the leave-out test set of TCGA-BRCA cohort. (b) PR curve and AUC value achieved by our model on the leave-out test set of TCGA-BRCA cohort. (c) Visualization of attention heatmaps of each molecular subtype. The left column showed the original slides, and the middle column showed the heatmaps generated by coloring each patch according to its learned attention score. The right column showed some representative patches assigned highest attention scores of each molecular subtype.

### 2.7 Computational pathological feature established an independent prognostic factor

We developed a risk model to assess patient prognosis using a Cox proportional hazards deep neural network [28]. The model incorporated slide-level pathological image features, gene expressions, clinical data, PAM50 molecular subtypes, and other potential factors. We constructed the prognosis risk model using negative log-likelihood as the loss function and predicted overall survival and progression-free interval. The predictive performance was evaluated using the average concordance index (C-Index) on three-fold cross-validation. Kaplan-Meier curves were used to visualize the stratification quality between low-risk and high-risk patients. In the experiment with overall survival as the endpoint, our model achieved a concordance index of 0.685 on the test set. Figure 6(a) showed that the high-risk group had a significantly lower survival time than the low-risk group (p*<*0.001). A univariate Cox regression analysis of the predicted risk score and other clinical data was conducted and displayed in a forest plot in Figure 6(b). The risk score based on pathological features was found to be a significant independent factor affecting overall survival (HR=3.205, p*<*0.001). In the experiment with progression-free interval as the endpoint, our model achieved a concordance index of 0.647 on the test set. Figures 6(c-d) illustrated that the high-risk group had a significantly higher prognosis risk than the low-risk group (p*<*0.001), and a univariate Cox regression analysis indicated that the risk score constructed by pathological features was a significant independent factor affecting progression-free interval (HR=4.695, p*<*0.001).

**Fig. 6:**
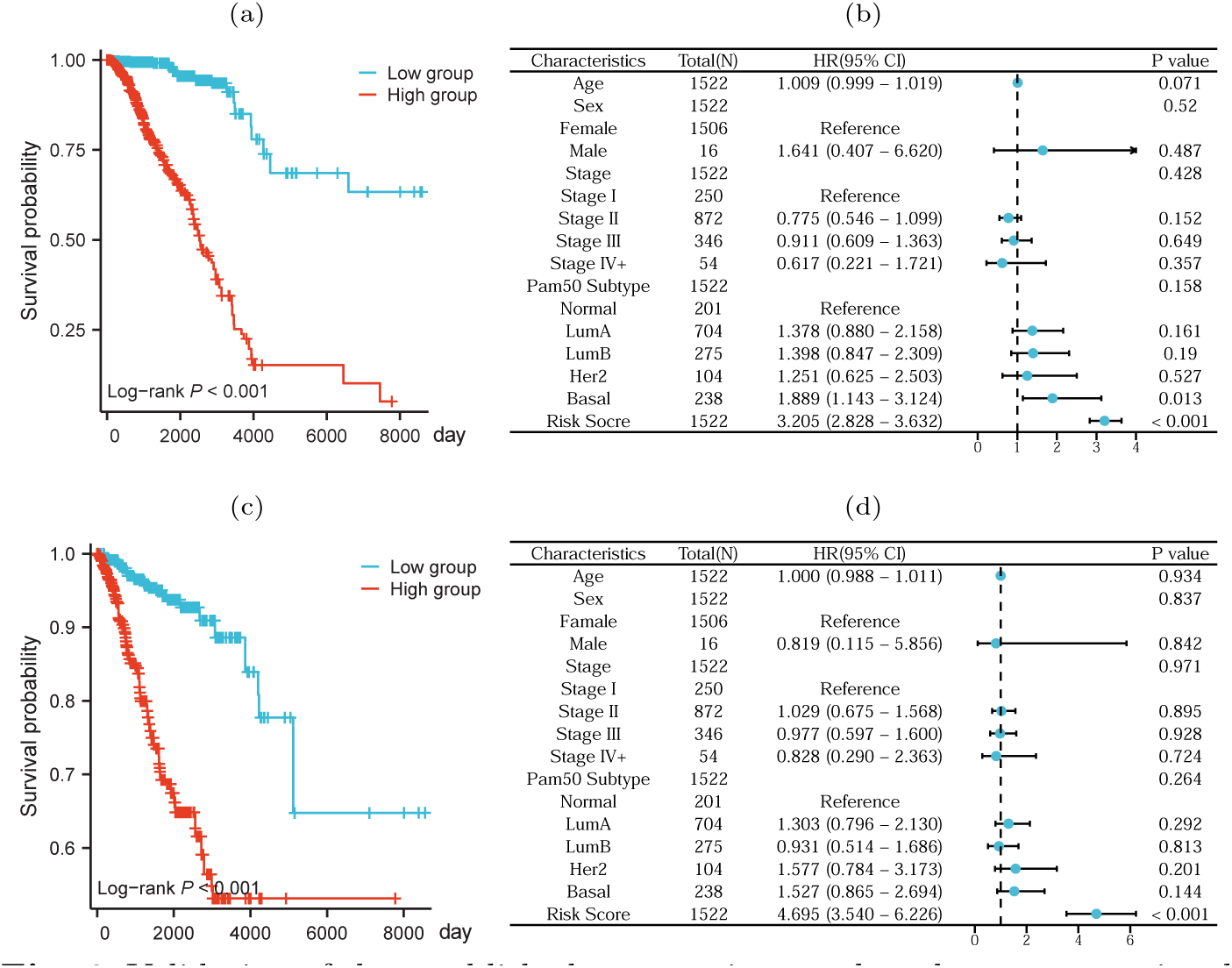
Validation of the established prognostic score based on computational pathological feature. (a) and (c) showed the Kaplan-Meier survival curves (Pvalue*<*0.05) of the lowand high-risk breast cancer patients stratified by the calculated prognostic scores for the endpoint event of overall survival and progression-free interval. (b) and (d) showed the forest plots of univariate COX regression regarding overall survival and progression-free interval on the TCGA-BRCA cohort. The forest plot showed the hazard ratios and confidence intervals of each factor, including the prognostic score, age, pathological stages, and molecular subtypes.

### 2.8 Model ablation showed contrastive learning and attentive pooling improve performance

To validate that the features extracted through contrastive learning-based pretraining enhance the performance of downstream tasks, we conducted a comparative analysis of two contrastive learning algorithms AdCo [29] and MoCo v2 [30], using ResNet50 [31] as the benchmark feature extractor. Our evaluation metrics included the Spearman correlation coefficient and Pearson correlation coefficient [29]. Figure 7(a) illustrated the performance of computational pathological features extracted by different algorithms in predicting gene expression levels. The experimental results indicated that the features extracted by AdCo contrastive learning yielded the most optimal performance, with an average Pearson and Spearman correlation coefficient of 0.41 and 0.37, respectively. Furthermore, we observed that both contrastive learning-based features significantly outperformed ResNet50 in this task.

**Fig. 7:**
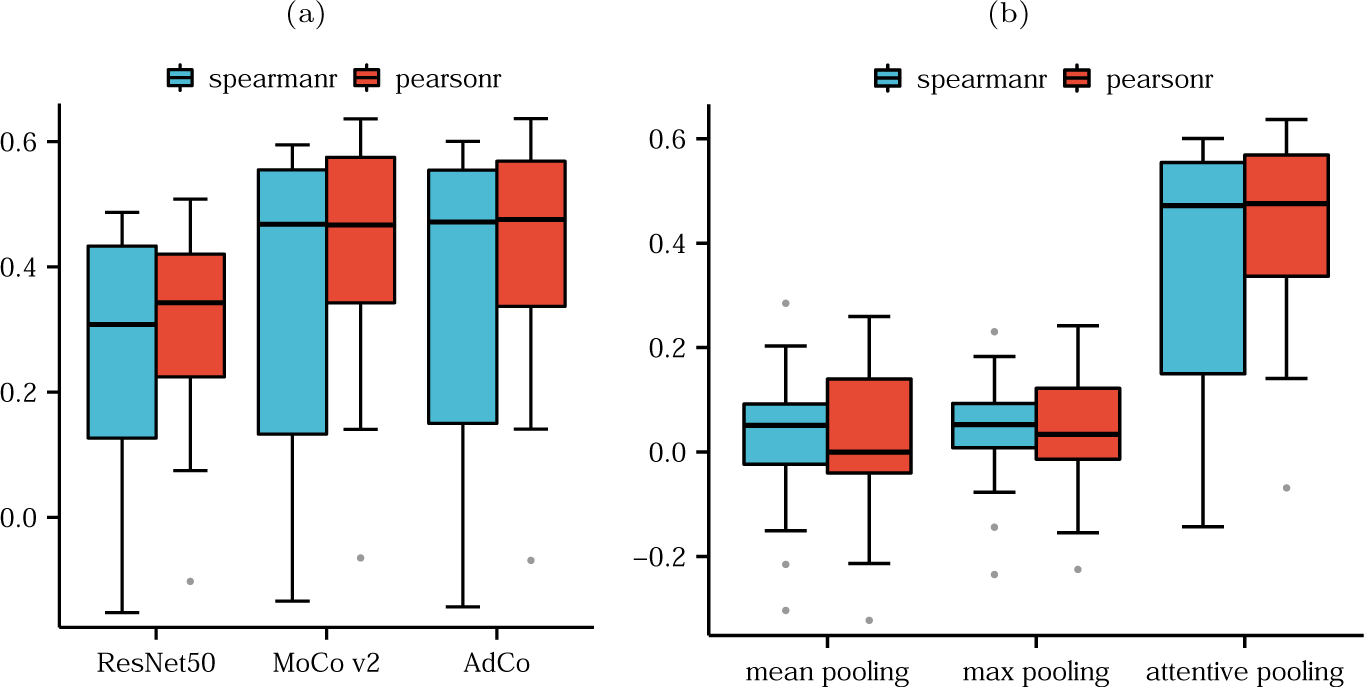
Model ablation verified that contrastive learning and attentive pooling improve performance. (a) Boxplots of the correlation coefficients achieved by ResNet50 baseline and two contrastive learning methods (MoCo v2 and AdCo) in predicting the gene expression levels of 21 recurrence-related genes in breast cancer. (b) Boxplots of the correlation coefficients obtained by different feature aggregation strategies, including mean pooling, max pooling, and attention pooling.

We also examined the influence of various feature aggregation strategies on predictive performance by testing max pooling, average pooling [30], and attentive pooling on gene expression levels. As depicted in Figure 7(b), attentive pooling achieved performance far superior to the other two feature aggregation methods.

### 2.9 Spatial transcriptomic data validated spatial deconvolution of gene expressions

Spatial transcriptomics combines microscopic imaging and RNA sequencing technology to capture gene expression data while maintaining the spatial location of cells within the tissue. This allows the mapping of RNA-seq data onto tissue sections, enhancing the comprehension of the spatial organization of cells and their gene expression patterns [32]. To verify our model capacity to perform spatial deconvolution, we produced heatmaps colored by patch-level attention scores of two biomarker genes, MKI67 and BCL2. Meanwhile, we obtained spatial transcriptomics data from 10x Genomics and illustrated the spatial landscapes of these two gene expressions on the pathological image. As shown in Figure 8(a-b), we observed that the attention heatmaps of MKI67 and BCL2 were highly consistent with their spatial expression landscapes derived from the spatial transcriptomics.

**Fig. 8:**
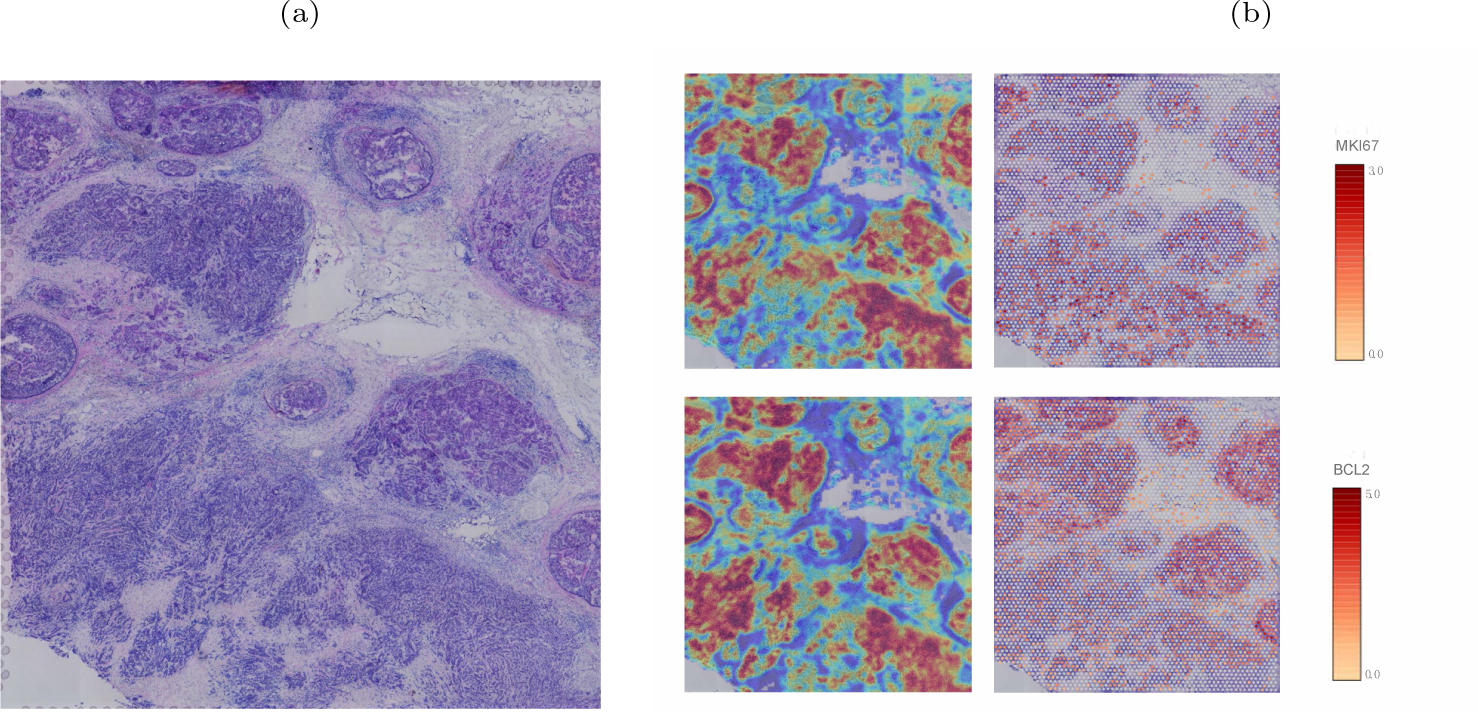
Visual validation of our model ability to spatially deconvolute the gene expression using spatial transcriptomic data. (a) Original pathological section used for spatial transcriptome sequencing. (b) Heatmaps generated by our model and spatial transcriptomic data showing the spatially expressed landscapes of the MKI67 and BCL2 genes.

### 2.10 Effective prediction of trastuzumab treatment outcome

Our final analysis evaluated whether our model can capture pathological image features to predict the treatment response of breast cancer patients to trastuzumab. We obtained a cohort of patients treated with trastuzumab from Yale University. This cohort included the patients with a pre-treatment breast core biopsy with HER2 positive invasive breast carcinoma who then received neoadjuvant targeted therapy with trastuzumab. A case was designated as responder if the pathological examination of surgical resection specimens did not report residual invasive, lympho-vascular invasion or metastatic carcinoma, and otherwise non-responders. After removal of the H&E images without enough detectable foreground tissue by redefined area threshold, we obtained 75 cases (34 responders and 41 non-responders).

Given that trastuzumab is a monoclonal antibody targeting HER2, we fine-tuned the feature encoder using HER2 level measured by Reverse-Phase Protein Array (RPPA) assay obtained from TCGA. This enabled our model to capture morphological features indicative of HER2 expression from pathological images. We employed five-fold cross-validation and used the ROC-AUC to evaluate model performance. Our results, depicted in Figure 9(a), demonstrated that our model achieved an AUC of 0.72 (95% CI: 0.65-0.81). In addition, we compared heatmaps generated by our model with tumor regions annotated by pathologists at Yale University and observed a high degree of consistency between the them, as shown in Figure 9(b). These findings support the feasibility of using pathological images to predict treatment response to trastuzumab, and demonstrate the effectiveness of deep learning models in identifying tissue morphological features associated with specific drug responses.

**Fig. 9:**
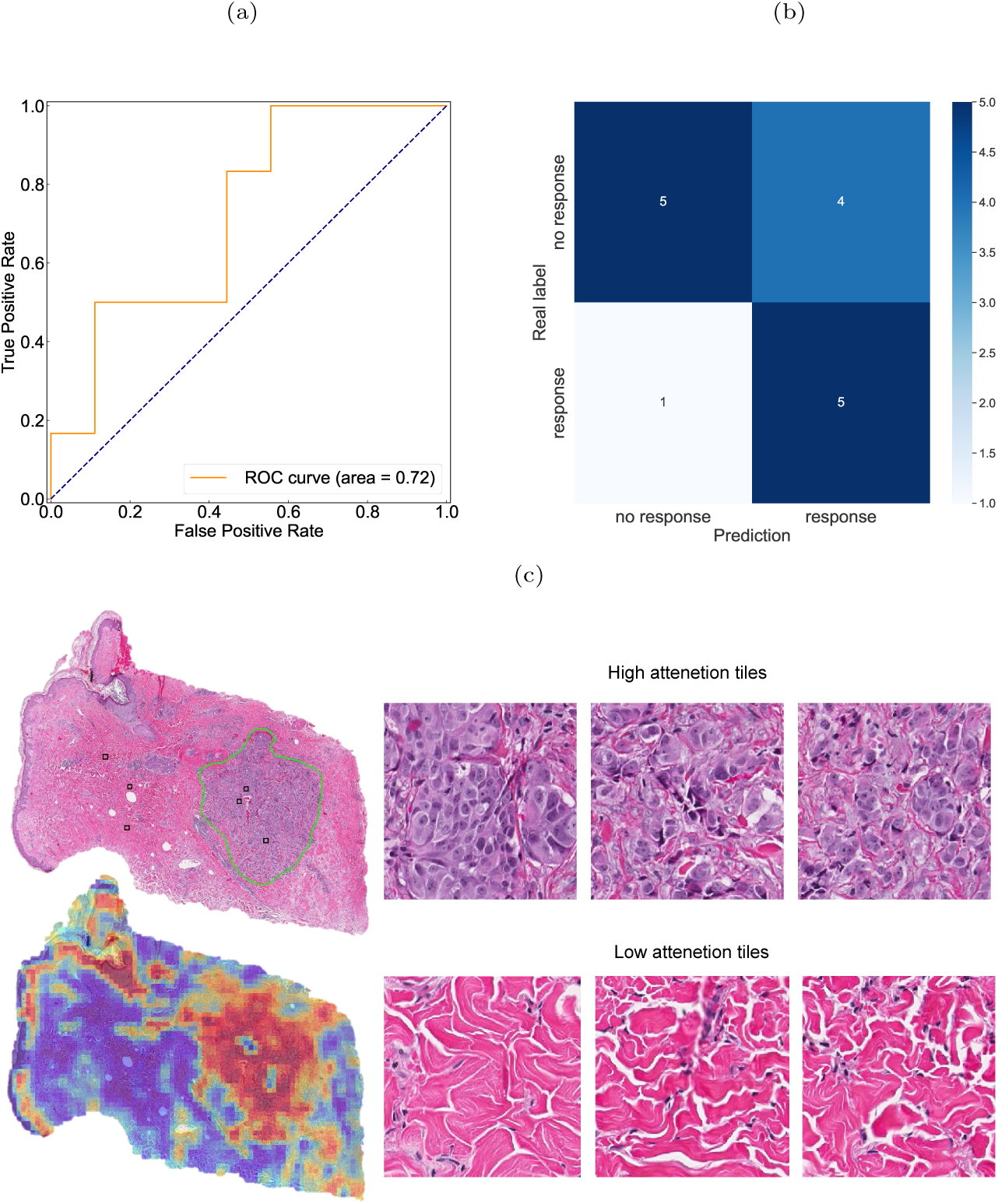
Performance evaluation of our model in inferring drug response on the Yale trastuzumab response cohort of breast cancer. (a-b) ROC curve and confusion matrix achieved by our model in predicting the drug response (c) Attention heatmap generated by coloring each patch by its attention score learned in the prediction of trastuzumab response. The highlight regions rendered in red and blue color were crucial in predicting drug response. The left column showed the tumor regions annotated by a pathologist and heatmap generated our model. The right column shows some representative patches assigned high and low attention scores representing their importance for drug response.

## 3 Materials and methods

### 3.1 Whole Slide Image

Whole slide images stained with H&E provide a detailed view of cellular components, allowing for easy differentiation between cell nuclei and cytoplasm. We obtained breast cancer whole slide images from TCGA-BRCA cohort for model training. The TCGA-BRCA cohort contains 1,979 fresh frozen WSIs and 1,131 formalin fixed paraffin-embedded (FFPE) WSIs from 1,094 breast cancer patients. The slide-level diagnosis provided by TCGA database were used as ground truth for classification labels. The PAM50 molecular subtyping was also obtained from the TCGA database.

Three other independent cohort were included for model evaluation. The first was the CPTAC-BRCA cohort that includes 642 fresh frozen WSIs from 134 patients of breast cancer. It was used to validate our model performance to infer gene expression levels using extracted pathological image features.

The second independent cohort came from the Changzhou Second People’s Hospital (CZSPH) in Jiangsu, China. The CZSPH cohort collected 91 FFPE WSIs from 90 breast cancer patients, including 15 recurrence cases within 5 years. The CZSPH cohort was used as an external validation set for recurrence prediction.

The third is the Yale trastuzumab response cohort that was used to evaluate model ability in predicting drug response. The Yale cohort contained 75 FFPE WSIs of 75 cases of breast cancer (34 responders and 41 non-responders). A case was designated as responder if the pathological examination of surgical resection specimens did not report residual invasive, lympho-vascular invasion or metastatic carcinoma, and otherwise non-responders. Note that only the slides with a magnification greater than 20* were included in this study.

### 3.2 Slide annotation

A pathologist with more than ten-year experience was asked to manually annotate a few slides. The annotated slides were used for the validation of spatial localization of tumor and necrosis regions. During annotation, the pathologist was blinded with regard to any molecular or clinical feature.

### 3.3 RNA-seq dataset

We collected the RNA-seq data matched to whole slide images from TCGABRCA and CPTAC-BRCA cohorts to train and validate our model capability in predicting gene expression levels, respectively. We adopted the upper quartile normalized (FPKM-UQ) fragments per kilobase per million mapped fragments (FPKMs) as expected values to train the model. Since the range of gene expression levels spans several orders of magnitude, straightforward regression on the raw RNA-Seq data would cause the model to focus only on the most strongly expressed genes, neglecting the signals of other genes. To overcome this problem, we applied a log10(1+x) transformation to the gene FPKM-UQ expression values [17].

### 3.4 Spatial transcriptomic data

We acquired spatial transcriptomic data from a breast cancer specimen using the Visium Spatial Gene Expression protocol from 10x Genomics. To explore the spatial locations of gene expression, we utilized the 10x Loupe Browser, a desktop application that offers interactive visualization capabilities for various 10x Genomics solutions. This enabled us to visualize the spatial landscape of specific biomarker genes experssion across the pathology image.

### 3.5 Preprocessing of whole-slide images

Due to the ultra-high resolution of pathological images, it is inhibitive to directly use them as inputs of deep neural networks. Therefore, we tessellated WSIs into small-size squares (patches) so that they were ready as input of deep learning model. For each slide, we used OpenSlide to read it into memory and then convert RGB to the HSV color space. For tissue regions (foreground), we generated a binary mask by thresholding the saturation channel in the HSV color space, smoothing edges and performing morphological closure to fill small gaps and holes. This effectively segmented the WSI image into tissue and non-tissue regions. After segmentation, each WSI was split into 256×256 pixel patches within the foreground region and filter out white image patches without enough tissue (pixel background value *>*210). It is worth noting that in order to improve access efficiency, we saved only the coordinates and metadata of each patch in HDF5 format.

### 3.6 Contrastive learning for feature extraction

Due to the scarcity of pixel-level labels of pathological images, traditional fully supervised learning is not applicable in our study. We used self-supervised contrastive learning on large-scale unlabeled patches to learn latent representations. In our study, we adopted the Adversarial Contrastive learning (AdCo) algorithm for pretraining. AdCo stands for adversarial contrastive for efficient learning of unsupervised representations from self-trained negative adversaries. It relies on constructing a collection of negative examples that are sufficiently hard to discriminate against positive queries when their representations are self-trained. We also tested another contrastive learning algorithm, MoCo v2, which is an improved version of the Momentum Contrast self-supervised learning algorithm. A ResNet50 network was used to benchmark the performance of contrastive learning.

For contrastive learning, the positive pairs were generated through data augmentation techniques such as image rotation, random cropping, and color transformation. Formally, denote the input patch as by *x_i_*, we used the ResNet50 CNN network as the backbone encoder, assuming that the encoder f transforms the patch into a latent representation *h_i_* = *Z*(*x_i_*), which was then projected onto an embedding *q_i_*through multiple fully-connected layers. A negative sample queue was initialized to store the latent representations of the patches. Contrastive learning trained the network parameters *θ* to distinguish between the query sample *q_i_* and its positive partner from a set of negative samples K. The contrastive loss is defined as:

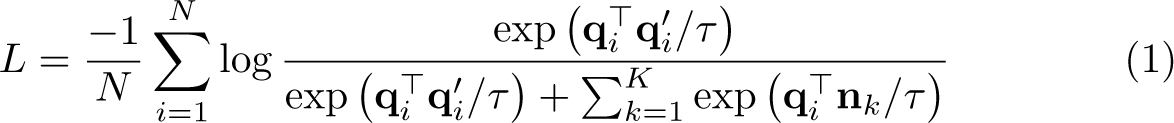

where *q_i_* and *q_i_* represent the embeddings of a pair of positive samples, *n_k_* refers to the embedding of a negative sample and *τ* is the temperature parameter.

Inspired by adversarial learning, AdCo aimed to generate challenging negative samples to be distinguished from query samples. It extended the memory bank that was used to stack negative sample embeddings by regarding negative samples as learnable weights. This allowed the learned negative samples to closely track changes in the learned embedding representations. AdCo taked the embedding of all samples as negative adversaries to network *θ* and updated the negative samples to maximize the contrastive loss, so that the adversarial negative samples were push closer to query samples. So, we haved the following min-max problem:

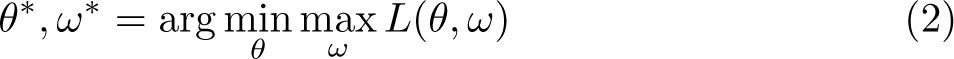

in which *ω* represents the set of dynamically updated adversarial negative samples. In fact, *ω* can also be considered as a set of model parameters. During the training process, the network parameter *θ* was updated along the direction of gradient descent, while *ω* was updated along the direction of gradient ascent.

Thus, the parameters *θ* and *ω* were alternately updated as follows:

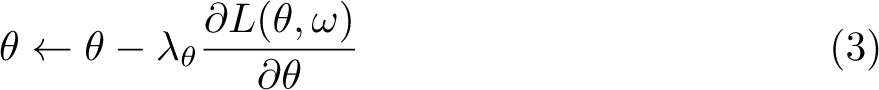

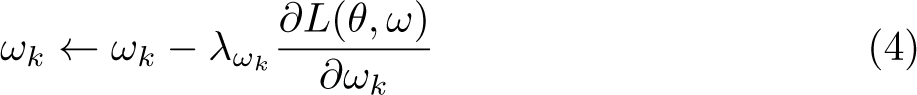

Once the contrastive learning-based pretraining was completed, we obtained the low-dimensional embedding of each patch using the trained ResNet50 backbone encoder. As a result, we converted each 256*256px patch into a 1024-dimensional vector. The extracted features were stored in HDF5 format ready for downstream tasks.

### 3.7 Attentive pooling for patch-level feature aggregation

Since the downstream task was based on slide-level features, we aggregated patch-level features into slide-level features. Unlike max pooling or average pooling, we used gated attention pooling here. Suppose a slide is divided into n patches S=*{p*_1_*, p*_2_*, …p_n_}*, the gated attention pooling is essentially a weighted average pooling where the weight is trainable.

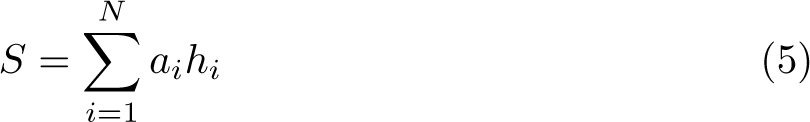

in which *a_i_* is the learnable parameter indicating the importance of *i*-th patch to specific task. Formally, *a_i_* is defined as below:

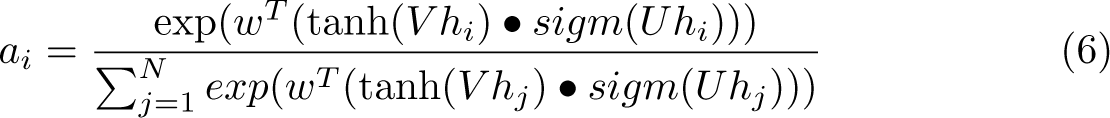

where *U* and *V* are all learnable parameters, *•* is an element-wise multiplication and *sigm*() is the sigmoid activation function. and *w^T^*is used to transfer the feature vector into raw attention score. The major difference between gated-attention pooling and max/mean pooling was that it introduced learnable parameters that were iteratively updated during the training process, making the model highly flexible and adaptable to different downstream tasks. More importantly, the attention weights after training reflected the importance of patch-level features to downstream task, which makes the model interpretable.

### 3.8 Model training and inference

During the contrastive learning phase, we randomly selected a fraction of patches from each slide and stacked them in the memory bank, since the large number of patches exceeds the memory capacity. This is a reasonable approach because individual patches have relatively low information content compared to natural images, so that a certain proportion of randomly extracted patches enables us to obtain the overall information of the slide. The adversarial negative samples were dynamically updated to force the encoder to capture the essential features of each patch and distinguish it from others.

In our implementation, ResNet50 was used as the backbone network. We only used its first four layers (the output of the last layer is 1024) and loaded the weights pre-trained on ImageNet. Two fully connected layers were appended to the model and used to project the representation into the 128-dimensional latent space[33]. SGD[34] was used as the optimizer. The learning rate of the backbone network was set to 0.03. Weight decay was set to 0.0001 and momentum was set to 0.9.

### 3.9 Downstream tasks

We applied transfer learning to leverage slide-level features for various downstream tasks, such as tumor diagnosis, gene expression level estimation, PAM50 molecular subtyping, recurrence and drug response prediction.

We performed tumor diagnosis on the TCGA-BRCA cohort, using the slide label (tumor or non-tumor) provided by the TCGA database as the ground truth. We employed a fully-connected layer plus a softmax layer as the prediction model that took the slide feature as input. The cross-entropy was used as the loss function for the classification task. Denote by *y* and *y^′^* the true and predicted label for tumor diagnosis, the loss function is as below:

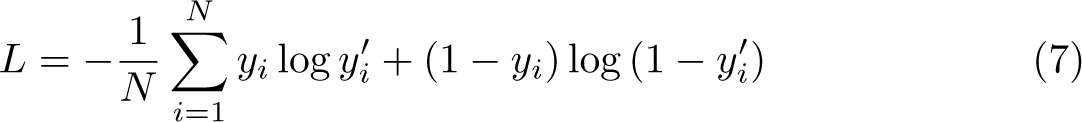

The gene expression level estimation was formulated as a regression task. Taking as input the extracted slide features, we used a multi-task learning model with a fully-connected layer and an output layer to predict the gene expression level. For instance, the output layer had 21 nodes corresponding to the expression levels of 21 genes associated with breast cancer recurrence. In our practice, we have test multiple hidden layers, and found that a single hidden layer could achieve superior performance in the regression task. We used the mean squared error as the loss function:

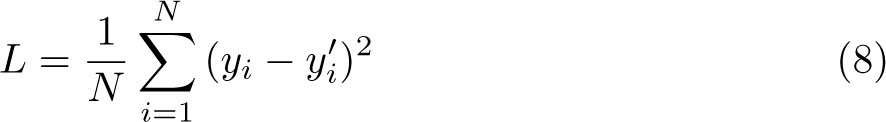

For PAM50 molecular subtyping, it was formulated as a multi-class classification problem. The model consisted of a fully-connected layer and a softmax layer. Because the normal-like subtype is possible to be more likely caused by contamination of normal breast tissue present in the specimen. We excluded the normal-like subtype in the molecular subtype prediction task. As a result, the output layer included four nodes corresponding to four molecular subtypes, namely luminal A, luminal B, HER2-enriched, and basal-like.

For recurrence prediction, a case was labeled as 1 if it had recurrence, metastasis, or a new tumor event within five years, and 0 otherwise. For drug response, a case was designated as responder if the pathological examination of surgical resection specimens did not report residual invasive, lympho-vascular invasion or metastatic carcinoma, and otherwise non-responders. Therefore, the prediction of recurrence and drug response from pathological features were both formulated as binary classification problems, and the cross-entropy loss function was used.

During the training process, we used the Adam[35] optimizer with a weight decay of 0.0001. For tumor diagnosis, gene expression level estimation, PAM50 molecular subtyping and recurrence prediction, we applied a decay strategy to dynamically adjust the learning rate. The learning rate was multiplied by 0.1 every time the model trained for 10 epochs. For drug response prediction, we used a cosine annealing function to dynamically adjust the learning rate.

### 3.10 Establishment of prognosis risk model

To build a prognosis risk model based on slide-level features, we employed a deep learning-based Cox proportional hazard regression model. The formula of the risk function is as follows:

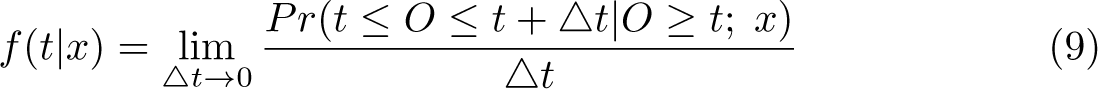

It estimates the instantaneous death rate of individual *x* at time *t*. The Cox proportional hazard regression models it as:

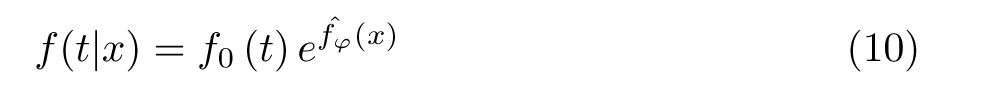

where *x* is the input feature that represents the baseline risk function at time *t*. In this study, we concatenate the slide features, clinical data, and the 21 gene expression levels into a one-dimensional vector as the model input. We built a multi-layer perceptron model composed of five fully-connected layers. The network output was a single node using the SELU activation function, which estimated the log hazard function in the COX model. We trained the network by setting the objective function to the regularized average negative log partial likelihood as below:

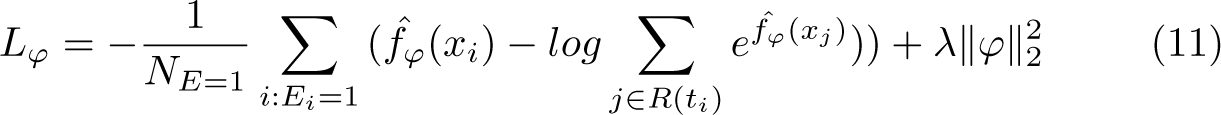

In which *R*(*t_i_*) is the risk set at time *t_i_* indicating the set of patients who are at risk at time *t*, *E_i_*= 1 indicates that the observed event (death) occurred, *N_E_*_=1_ is the number of samples with observed events, and *λ* is the *ℓ*_2_ regularization parameter. During the training process, Adam was used as the optimizer, the learning rate was set to 0.01, the weight decay was set to 0.001 and the regularization parameter was set to 0.15.

## 4 Discussion and Conlusion

Genetic testing has been an important clinical examination for tumor subtyping and targeted drug delivery. However, due to technology and cost reasons, genetic testing has not become clinically spread, especially in developing countries. The rapid advancement of digital pathology motivated the application of computational pathology to infer genetic mutations, microsatellite instability and tumor microenvironment. Following the rationale that the change of cell and tissue phenotype is driven by the variation of gene expression pattern, we have explored the computational pathology-based gene expression prediction for breast cancer. Inference from pathological features to molecular patterns relies heavily on the powerful feature extraction capabilities of deep learning, while self-supervised contrast learning performs well in representation learning but has not yet been exploited in digital pathology mining. Our experiments demonstrated that pretraining based on contrast learning can extract better representation embeddings and effectively improve the performance of molecular-level feature prediction. And the attention mechanism assigns trainable weights to each patch based on weakly supervised signals, reflecting the importance of molecular expression levels within different regions of the tumor. This not only explored model interpretability but also identified morphological features at different molecular expression levels. Although current pathology image-based approaches are not yet up to the standard of clinical application, we believe that with the accumulation of data, especially with the development of spatial transcriptomics and single-cell histology, new computational pathology-based approaches will emerge to predict molecular features from a multi-modal perspective.

In this paper, we proposed a neural network model based on contrast learning and attention mechanism for downstream tasks for tumor diagnosis, gene expression prediction, molecular subtyping and recurrence risk prediction on whole pathology images. Our experimental results showed that our model achieved good performance on tumor diagnosis tasks, molecular subtyping, recurrence prediction task, and drug response prediction task and that prognostic scores constructed from computed histopathological features can be used as independent prognostic factors. Notably, our model was able to significantly predict gene expression levels from pathological images because multiple gene prediction tasks are learned together, sharing with each other through a shared representation at a shallow level, complementing each other’s learning to domain-relevant information, facilitating each other’s learning and enhancing generalization. In addition, our visualization of patches-level attention scores generated according to the attention mechanism is highly consistent with pathology annotations and spatial transcriptomes. We concluded that introducing contrast learning into the feature extraction stage greatly improved the problem of no effective annotation of pathology in the pathology image domain, and significantly improved a range of downstream tasks. In the future, we hope to use more contrast learning frameworks to fuse different multi-scale features and multi-modal features to effectively improve our models and to test our approach on more tumor datasets. In conclusion, we hope that our research and methods will bring new perspectives and contribute to the development of computational pathology.

## Declarations

- Funding This work was supported by National Natural Science Foundation of China (62072058, 82073339).
- Conflict of interest The authors declare no competing interests.
- Ethics approval
- Data availability The TCGA-BRCA diagnostic and frozen slides with corresponding labels, and the matched RNA-seq data are available from the NIH genomic data commons (https://portal.gdc.cancer. gov). The CPTAC-BRCA whole-slide with corresponding labels, and the matched RNA-seq data are available from the NIH cancer imaging archive (https://cancerimagingarchive.net/ datascope/cptac). The whole slide images and drug response data from Yale trastuzumab response cohort are available at TCIA database https://wiki.cancerimagingarchive.net/. The spatial transcriptomic data of a breast cancer specimen from 10X genomics are available at https://www.10xgenomics.com/. The All reasonable requests for academic use of in-house raw and analysed data can be addressed to the corresponding author. The WSIs of the CZSPH cohort, together with clinical data, are available at https://www.zenodo.org/deposit/7839324.
- Code availability The source and data are available at https://github.com/hliulab/wsi2recurrence.
- Authors’ contributions H.L., Y.Z. and A.Z. conceived the study and designed the experiments. Y.Z. performed the experimental analysis. Z.S and J.L. curated the in-house datasets and collected smartphone microscopy data. H.L. and Y.Z. prepared the manuscript. H.L. and J.L. jointly supervised the research.

